# Molecular modeling of apoE in complexes with Alzheimer’s amyloid-β fibrils from human brain suggests a structural basis for apolipoprotein co-deposition with amyloids

**DOI:** 10.1101/2023.08.04.551703

**Authors:** Emily Lewkowicz, Mari N. Nakamura, Michael J. Rynkiewicz, Olga Gursky

## Abstract

Apolipoproteins co-deposit with amyloids, yet apolipoprotein-amyloid interactions are enigmatic. To understand how apoE interacts with Alzheimer’s amyloid-*β* (A*β*) peptide in fibrillary deposits, the NMR structure of full-length human apoE was docked to four structures of patient-derived A*β*_1-40_ and A*β*_1-42_ fibrils determined previously using cryo-electron microscopy or solid-state NMR. Similar docking was done using the NMR structure of human apoC-III. In all complexes, conformational changes in apolipoproteins were required to expose large hydrophobic faces of their amphipathic *α*-helices for sub-stoichiometric binding to hydrophobic surfaces on sides or ends of fibrils. Basic residues flanking the hydrophobic helical faces in apolipoproteins interacted favorably with acidic residue ladders in some amyloid polymorphs. Molecular dynamics simulations of selected apoE-fibril complexes confirmed their stability. Amyloid binding *via* cryptic sites, which became available upon opening of flexibly linked apolipoprotein *α*-helices, resembled apolipoprotein-lipid binding. This mechanism probably extends to other apolipoprotein-amyloid interactions. Apolipoprotein binding alongside fibrils could interfere with fibril fragmentation and secondary nucleation, while binding at the fibril ends could halt amyloid elongation and dissolution in a polymorph-specific manner. The proposed mechanism is supported by extensive prior experimental evidence and helps reconcile disparate reports on apoE’s role in A*β* aggregation. Furthermore, apoE domain opening and direct interaction of Arg/Cys158 with amyloid potentially contributes to isoform-specific effects in Alzheimer’s disease. In summary, current modeling supported by prior experimental studies suggests similar mechanisms for apolipoprotein-amyloid and apolipoprotein-lipid interactions; explains why apolipoproteins co-deposit with amyloids; and helps reconcile conflicting reports on the chaperone-like apoE action in A*β* aggregation.

## INTRODUCTION

In amyloid diseases, normally soluble proteins and peptides form insoluble fibrils, causing cell and organ damage. Nearly 40 human proteins form pathologic amyloids either in the brain or in other organs, leading to various diseases [1]. Amyloid deposition in the brain is a hallmark of neurodegenerative disorders such as Alzheimer’s disease (AD), a debilitating incurable disease that affects millions of patients worldwide. Extracellular deposition of amyloid-beta (A*β*) peptide plays a key pathologic role at early stages of AD, while tau protein deposition is central to AD pathology at later stages [2–4]. Emerging therapeutic strategies for AD target A*β* amyloid [5].

Amyloid deposits contain the major fibril-forming protein, lipids, and other non-fibrillary components including apolipoproteins (apo), serum amyloid protein, and heparan sulfate (HS) proteoglycans [1, 6, 7]. These “amyloid signature proteins” provide diagnostic markers of amyloid and, in case of apoE and HS, its important modulators. Determining the molecular basis for amyloid co-deposition with these accessory molecules is necessary to understand their action and harness it for therapeutic applications[6, 8, 9].

New insights are emerging from the atomic structures of amyloid fibrils determined using cryo-electron microscopy (cryo-EM) and solid-state nuclear magnetic resonance (NMR) ([10, 11] and references therein). In amyloid structures, well-ordered cores contain stacks of flattened protein molecules forming parallel in-register intermolecular *β*-sheets (PRIBS), sometimes flanked by disordered “fuzzy coat” regions. In PRIBS, arrays of identical residues spaced at 4.7 Å run along the fibril sides. Depending on their composition, such residue arrays are either energetically favorable (e.g. polar ladders) or unfavorable (side chains with unbalanced charge or hydrophobicity). Charged residue arrays in amyloids can be stabilized by binding cofactors with complementary charge and geometry. Uncompensated basic arrays in amyloids have been predicted [12] and observed to bind periodic polyanions such as HS [13] or RNA [14]; conversely, uncompensated acidic arrays in amyloid can be stabilized by binding metal ions [15] or protons [12, 16]. The present study addresses stabilization of exposed hydrophobic residue arrays in amyloid.

Amyloid cores are enriched in hydrophobic residues that are exposed at the fibril ends or along the fibril side [11], potentially forming contiguous surfaces. Since solvent-exposed hydrophobic surfaces are energetically unfavorable and can trigger immune response [17], they are probably shielded *in situ* by amphipathic molecules, such as apolipoproteins and lipids that co-deposit with amyloids. Shielding of hydrophobic amyloid surfaces by anionic lipid micelles was observed in recent cryo-EM structures of recombinant *α*-synuclein fibrils [18]. However, no structures of apolipoprotein-amyloid fibril complexes are currently available. The present study proposes the first structural models of apoE-A*β* fibril complexes.

ApoE is a 299-residue glycoprotein that transports lipids in plasma and cerebrospinal fluid. ApoE secreted by hepatocytes circulates in plasma on high-density and very low-density lipoproteins and acts as a ligand for the receptor-mediated cellular uptake of very low-density lipoprotein remnants [3, 19, 20]. ApoE secreted by astrocytes and other brain cells forms high-density lipoproteins (diameter ∼10 nm) that transport lipids in the central nervous system and influence cellular uptake and clearance of A*β* [21]. ApoE is the major modulator of brain lipid metabolism and a critical player in AD and other neurodegenerative disorders [20, 22]; reduced levels of apoE are a causative risk factor for dementia [23].

Like other exchangeable (water-soluble) apolipoproteins, apoE is a dynamic molecule that binds reversibly to lipid surfaces *via* the large hydrophobic faces of its amphipathic *α*-helices [24]. Since these faces are sequestered in lipid-free apoE, lipid binding involves large conformational changes [25–28]. ApoE is comprised of two domains, a 22 kDa N-terminal (NT) domain (NTD) that contains the receptor and HS binding site, and a 12 kDa C-terminal (CT) domain (CTD) that forms the primary lipid binding site [3, 20, 26, 28, 29]. In lipid-free apoE, the NTD forms a four-helix bundle [29] while the intrinsically disordered CTD is involved in self-association [28]. On lipoproteins, both domains can acquire an extended predominantly *α*-helical conformation to directly bind lipids [28, 30]. Numerous biophysical studies using cross-linking and mass spectrometry, Förster resonance energy transfer (FRET), electron paramagnetic resonance (EPR) and other methods showed that lipid binding by apoE involves domain separation followed by the helix bundle opening in NTD to expose its hydrophobic helical faces to lipid [3, 19, 25–28, 30, 31]. Besides lipids, both apoE domains can bind A*β*, but the binding mechanism is unclear ([3, 8, 32–36] and references therein).

ApoE has been front and center in AD studies since 1990s when the apoE4 isoform emerged as the major genetic risk factor for the sporadic late-onset AD [37, 38]. The three major isoforms of human apoE differ at residues 112 and 158: C112 and C158 in apoE2, C112 and R158 apoE3, and R112 and R158 in apoE4. These residues confer subtle dynamic differences in NTD stability, which increases in the order apoE4 < apoE3 < apoE2 [39]; NTD-CTD interactions, which are weakest in apoE4 [25]; and susceptibility to proteolytic fragmentation, which is highest in apoE4 and possibly contributes to A*β* pathology [40–42]. ApoE4 circulates at lower levels compared to apoE2 or apoE3 in plasma and cerebrospinal fluid, and apoE post-translational modifications [43] and lipidation are also isoform-specific [20]. These differences impact apoE interactions with lipoprotein receptors, lipids, proteoglycans, and A*β,* with major implications for AD and cardiovascular disease [19, 20, 44].

Although the critical role of apoE in AD has been firmly established, the underlying mechanisms are far from clear. ApoE interacts with A*β* peptide in an isoform-specific manner and influences its homeostasis through several routes, including lipid transport, receptor-mediated A*β* clearance, and A*β* amyloid deposition [45]. The latter is thought to involve direct A*β*–apoE interactions [3, 8, 22, 46, 47]. However, the evidence from the animal model studies of the role of these interactions was conflicting (reviewed in [3, 8, 22, 47, 48]). Some mouse model studies reported that overexpression of murine or human apoE promotes A*β* deposition [49] and accelerates A*β* amyloid seeding, with apoE4 enhancing seeding more than apoE3 [50]. Others reported that apoE delays A*β* deposition, with apoE4 showing the smallest and apoE2 the largest effect ([48, 51] and references therein). These discrepancies were proposed to stem, in part, from the difficulty in generating mouse models relevant to specific aspects of human AD ([22, 48] and references therein).

*In vitro* studies of apoE-A*β* interactions were also contradictory [8, 52, 53]. Depending on the exact conditions, apoE has been proposed to act either as a “pathologic chaperonin” promoting A*β* aggregation [8] or as a beneficial anti-amyloid chaperonin [53]. Still, the consensus is that apoE binds with tens of nM affinity to *β*-sheet-rich A*β* aggregates and fibrils, thereby stabilizing them and interfering with amyloid formation [33, 35, 52–56]. Numerous studies showed that amyloid formation by A*β* enhances its binding to apoE, which involves multivalent interactions with both apoE NTD and CTD ([32, 35, 36, 47, 55, 57] and references therein). Conversely, apoE binding influences A*β* aggregation *in vitro*, although the results vary depending on the protein source and concentration, apoE post-translational modifications, lipidation, and other factors that alter the complex time-dependent patterns of self-aggregation and co-aggregation of both proteins [22, 52, 53, 55]. Earlier studies reported that apoE accelerates A*β* aggregation, with apoE4 having the strongest effect [36, 57, 58]. Subsequent studies using multiple biophysical, biochemical and cell-based approaches [35, 52, 54, 55, 59, 60] reported that at low micromolar protein concentrations [52] that approximate *in vivo* conditions [53], apoE delayed A*β* amyloid formation. Specifically, apoE delayed A*β* amyloid nucleation [52, 53, 55, 60, 61] and/or impaired fibril elongation and maturation ([52, 53, 55, 56, 62] and references therein). Unraveling molecular underpinnings of these complex effects and the origins of their apparent discrepancies is paramount for their therapeutic targeting ([8, 22, 52] and references therein). The present study suggests that a previously unappreciated key factor in apoE-A*β* amyloid interactions is amyloid polymorphism.

This study explores the structural basis for apoE binding to A*β* fibrils. We harness previously determined cryo-EM structures of A*β*_1-40_ and A*β*_1-42_ fibrils isolated from brains of patients with AD and other neurodegenerative disorders [15, 63]. Additionally, we use the solid-state NMR structure of seeded recombinant A*β*_1-42_ fibril that mimics features of the amyloid seed extracted from human AD brains [64]. By docking to these fibril structures the solution NMR structure of modified human lipid-free apoE3 [28], we propose the first molecular models of A*β* fibrils in complexes with full-length apoE. Molecular dynamics (MD) simulations of selected apoE-A*β*_1-42_ fibril complexes demonstrated their stability. Prior experimental studies of apoE-A*β* interactions verified key aspects of the proposed models. Additional support comes from docking the NMR structure of apoC-III (79 a. a.) [65], which is an A*β*-binding apolipoprotein and a biomarker of AD [66] with apparent neuroprotective properties [67]. The results reveal the driving forces for apolipoprotein binding to amyloid fibrils, help reconcile conflicting evidence from prior experimental studies, explain why apolipoproteins co-deposit with amyloids, and provide a structural basis for understanding how apolipoproteins can modulate amyloid formation, proliferation and interactions with other factors.

## METHODS

### Atomic structures of human Aβ fibrils and apolipoproteins E and C-III

To model apoE-A*β* fibril complexes, we used atomic structures of four different A*β* fibrils representing human amyloids (supplemental Fig. 1). These include three cryo-EM structures of fibrils isolated *post mortem* from patents’ brains. Fibrils of A*β*_1-40_ morphology I (4.5 Å resolution, PDB ID: 6SHS) were from vascular deposits in AD brain. Fibril structures of A*β*_1-42_ were reported in two morphologies, type I (associated with sporadic and familial human AD) and type II (found in other human neurodegenerative diseases and in a mouse model of AD) [15]. The cryo-EM structures for type I (sporadic AD, 2.5 Å resolution, PDB ID: 7Q4B) and type II (pathologic aging, 2.8 Å resolution, PDB ID: 7Q4M) were used for docking. Additionally, we used the solid-state NMR structure of recombinant human A*β*_1-40_ fibrils (PDB ID: 2M4J), which were grown *in vitro* from the seed isolated from AD brain tissue [64].

For apoE, we used the NMR solution structure of full-length human lipid-free apoE3 (PDB ID: 2L7B, supplemental Fig. 2a) [28]. To stabilize the flexible CTD and prevent apoE self-association, five CTD residues have been substituted to facilitate structural NMR studies (F257A/ W264R/ V269A/ L279Q/ V287E) [28]. Restoring these residues to their original composition in the apoE model caused no substantial changes in our docking results.

In addition, we used the NMR solution structure of full-length human apoC-III in complex with sodium dodecyl sulfate micelles (PDB ID: 2JQ3, supplemental Fig. 3a), which represents the lipid-bound conformation [65]. The structures of apoE and apoC-III were docked to the A*β* fibril structures as described below.

### Docking protocol

The published fibril structures, which contained 3-6 rungs, were elongated by adding duplicate molecular layers in Chimera using the measured symmetry to ensure that the fibril length exceeded the apolipoprotein length and provided ample space for docking. Docking of the full-length apolipoproteins and their fragments to the elongated fibril structures was carried out on the ClusPro Server (https://cluspro.org/login.php) [68]. The fragments were aligned along the fibril as described in Results and stitched together using COOT software [69]. The results were displayed using Chimera or VMD software. Docking of apoE2, apoE3 and apoE4 isoforms gave similar results that are reported for apoE4.

### Molecular dynamics (MD) simulations

MD simulations of the docked apoE-Aβ fibril complexes were carried out in NAMD version 2.13 [70]. The model was solvated in a water box in VMD using a cushion of 15 Å and the system was neutralized and brought to 0.15 M NaCl concentration using the Solvate and Ionize plugins in VMD [71]. The system was energy-minimized in two steps to remove initial poor interactions. First, 500 steps of conjugate gradient minimization were performed with all protein atoms fixed to relax the solvent structure. Then, a constrained minimization of the protein was carried out with harmonic constraints placed on the backbone atoms (force constant of 5 kcal/mol*·*Å^2^) and side chain atoms (force constant of 1 kcal/mol*·*Å^2^). The constraints were released slowly over a total of 5,000 steps. Next, the system was heated to 300 K at constant volume using harmonic constraints to the minimized coordinates (force constant of 1 kcal/mol*·*Å^2^). After reaching 300 K, the constraints were slowly removed over the next 500 ps of simulation time; the Langevin piston barostat was used to bring the pressure up to 1 atm. Production runs were carried out at 300 K, 1 atm for 100 ns using a final time step of 2 fs.

## RESULTS

### ApoE docking to Aβ_1-40_ fibrils from AD vasculature

*Ex vivo* fibril structures of A*β*_1-40_ were reported in three morphologies, I to III (Fig. 1a), containing one to three paired filaments with a right-handed twist packed side by side. Cryo-EM structures revealed a similar C-shaped flattened peptide in PRIBS conformation, with two dimer-forming molecules (residues 1-40) in the fibril core [63]. Morphology I structure, which was determined to the highest resolution, was used for docking. First, full-length apoE was docked to the fibril structure. This generated multiple unrelated docking poses around the fibril surface but no strong preference for binding at any specific site (supplemental Fig. 4a, top), suggesting that amyloid binding sites could be occluded in full-length lipid-free apoE. Next, apoE fragments were docked to A*β* fibrils. This strategy is widely used to identify cryptic sites that are hidden in apoproteins and require conformational changes for ligand binding [72]. The fragment-based approach is further justified by increased apoE fragmentation in AD, and by the observation that apoE fragments co-deposit with A*β* amyloid in human brain [40–42]. Initially, apoE was split into two fragments, NTD (1-165) and CTD (166-299) (supplemental Fig. 2b-c), which were docked separately (supplemental Fig. 4a, top). Of the 30 top docking poses predicted for each domain, almost all interacted with the fibril *via* molecular surfaces that are occluded in full-length protein. Therefore, domain separation in apoE can expose inner surfaces that are complementary to the fibril surface.

**Fig. 1.**
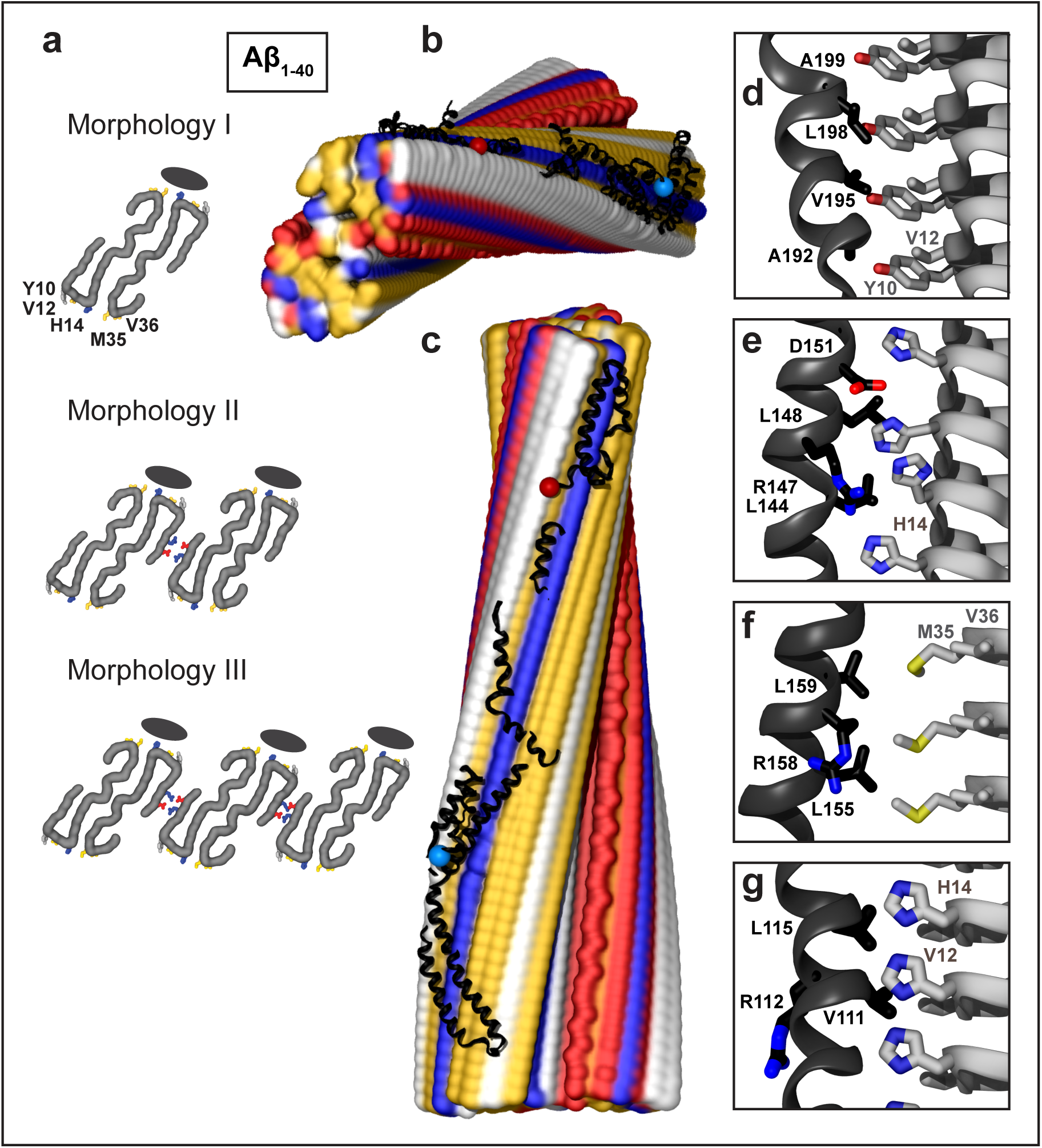
Structural model of apoE docked onto A*β*_1-40_ fibril from AD vasculature. The model was obtained by using apoE segments shown in supplemental Fig. 2d-h and the fibril structure (PDB ID: 6SHS). Segment positions are compatible with full-length apoE. (**a**) Fibril morphologies I-III containing one to three paired protofilaments (main chain in gray, view down the fibril axis). Morphology I was used for docking. Black ovals mark the apoE docking position at one of two predicted sites per paired protofilament. ApoE-coordinating residues Y10, V12, H14 from molecule 1 and M35, V36 from molecule 2 of A*β*_1-40_ protofilament are indicated. In morphologies II and III, residue pairs E3 and R5 that form salt bridges between adjacent protofilaments [63] are shown. (**b, c**) Top and side views of the docking model show apoE alongside A*β*_1-40_ protofilament. In this and other figures of apolipoprotein-amyloid complexes, apolipoprotein main chains are in black ribbons; blue and red dots mark N– and C-termini. Amyloid fibrils are in a surface representation: yellow – hydrophobic, white – polar, red – acidic, blue – basic (including His). Panels d-g show apoE-amyloid contacts within ∼5Å in selected regions (apoE – black, Aβ – gray). (**d**) A*β*_1-40_ residue ladder Y10, V12 forms hydrophobic interactions with apoE residues; A192, V195, L198, A199 from the apoE hinge region are shown. (**e**) H14 of A*β*_1-40_ forms mixed (polar/hydrophobic) interactions with apoE residues; R147, L144, L148, D151 from helix 4 are shown. (**f**) R158 in apoE3/E4, which flanks the hydrophobic face of helix 4, interacts unfavorably with M35 of A*β*_1-40_. (**g**) Hydrophobic residues of apoE helix 3 interact with residue ladders of V12 and H14 in Aβ_1-40_. Residue 112 (R112 in apoE4) points away from amyloid.

To probe if further exposure of apoE *α*-helices liberates their potential fibril-binding sites, the protein was divided into six fragments shown in supplemental Fig. 2d-h. The NTD was split into two pairs of antiparallel helices, a h1/h4 pair (residues 1-43 and 130-165, which were docked together to preserve their relative orientation) and a h2/h3 pair (residues 54-126). These helix pairings were chosen since they were shown to move away from each other during lipid binding ([25, 26, 28, 31] and references therein). The CTD was split into three helical fragments: the hinge (residues 166-203), lock (207-225) and CT region (238-299), which also move relative to each other upon lipid binding ([28, 31] and references therein). Disordered linkers (residues 44-53, 127-129, 204-206, and 226-237) were omitted from docking. Strikingly, all apoE fragments preferentially docked at the same hydrophobic site along the fibril (supplemental Fig. 5a, top). Each fragment docked with comparable likelihood in N-to-C or C-to-N orientation along the filament. By selecting proper orientations and translating the fragments along the fibril length, we were able to assemble them in a manner consistent with one full-length apoE molecule in a partially extended conformation.

Fig. 1 shows the resulting model. The hydrophobic faces of apoE *α*-helices interact with the A*β*_1-40_ residue ladders of Y10, V12 and H14 from molecule 1 and M35 and V36 from molecule 2 in the dimeric filament (Fig. 1a, d-f). While most apoE-amyloid interactions are hydrophobic (Fig. 1b-g), H14 rings form mixed hydrophobic and polar interactions (Fig. 1e). Such apoE binding is expected to stabilize amyloid for two reasons. First, the amphipathic *α*-helices of the bound apoE shield the most hydrophobic surface of amyloid from solvent. Second, bound apoE bridges two protofilaments in the dimeric A*β*_1-40_ filament, thus locking them together. Furthermore, apoE binding at this hydrophobic site is compatible with fibril morphologies II and III wherein the A*β*_1-40_ dimers or trimers pack laterally *via* charge-rich surfaces (Fig. 1a).

Notably, residues 112 and 158, which distinguish apoE isoforms 2, 3 and 4, are oriented differently in respect to amyloid: the 112 side chain points away from the fibril, while the 158 side chain, which flanks the hydrophobic helical face, projects towards the fibril and can interact with M35 of A*β*_1-40_ (Fig. 1f, g). Consequently, R158 in apoE3 and apoE4 interacts unfavorably with M35 on the fibril surface; the interaction is expected to become more favorable either upon M35 oxidation in A*β*_1-40_, which converts a hydrophobic Met into polar Met sulfoxide, or in apoE2 that contains C158 instead of R158. Furthermore, conformational opening of apoE domains, which is key to the proposed amyloid binding mode, is isoform-specific [25, 39].

In summary, apoE docking to the structure of A*β*_1-40_ fibrils from AD vasculature predicts apolipoprotein binding *via* its hydrophobic *α*-helical faces to the most hydrophobic surface alongside the fibril. The binding is expected to stabilize amyloid, is compatible with monomeric, dimeric and trimeric A*β*_1-40_ filaments, and the model suggests a structural basis for the observed apoE isoform-specificity.

### Overview and the docking strategy for Aβ_1-42_ fibrils from parenchymal deposits in AD

Cryo-EM structures of *ex vivo* A*β*_1-42_ fibrils were reported in several morphologies: type I and its dimeric version, type Ib (associated primarily with sporadic AD), and type II (associated with familial AD and other neurodegenerative disorders such as pathologic aging) [15]. All morphologies had a left-handed twist and contained two protofilaments comprised of S-shaped A*β* molecules in the amyloid core (residues 9-42 in type I and 11-42 in type II), with NT segments forming a “fuzzy coat”. Different protofilament packing generated different filament surfaces. In type I filaments, most hydrophobic residues were sequestered, leading to highly hydrophobic top and bottom surfaces (supplemental Fig. 1c). In type II filaments, more hydrophobic residues were exposed along the sides while the end surfaces were less hydrophobic (supplemental Fig. 1b). All filaments had a charge-rich surface, including an acidic ridge formed by E22 and D23 and stabilized by bound metal ions [15]. Since the origin of these unidentified surface-bound ions was unclear, they were not included in our docking models.

To avoid steric clashes between apoE and the “fuzzy coat”, the NT ends of the PRIBS core (residue 9 in type I and residue 11 in type II filaments) were blocked from docking. Otherwise, the docking strategy was similar to that for A*β*_1-40_. Briefly, full-length apoE showed no specific binding; splitting apoE into NTD and CTD fragments liberated helical surfaces that were better suited for amyloid binding (supplemental Fig. 4b-c). Further splitting into six apoE fragments (shown in supplemental Fig. 2d-h) produced consistent results for type II and type I fibrils as described below.

#### ApoE docking to type II Aβ_1-42_ fibrils and MD simulations

All apoE fragments showed a strong preference for docking at the same largely hydrophobic groove along the fibril side (supplemental Fig. 5b). By translating these fragments along the fibril length and selecting their relative N-to-C orientations, the fragments were combined into one full-length apoE molecule (Fig. 2a, b). The docking site for apoE was lined with A*β*_1-42_ side chains of V39 and I41 from one protofilament and E22, D23 and S26 from the other protofilament (Fig. 2c). The contacts involved van der Waals interactions between the hydrophobic helical faces of apoE and A*β*_1-42_ side chains of V39 and I41 (Fig. 2c, e, g). Additionally, the docking suggested favorable ionic interactions between the basic residue arrays that flank the hydrophobic helical faces in apoE and acidic ladders of D23 and, to a lesser extent, E22 in A*β*_1-42_ fibrils (Fig. 2c, f).

**Fig. 2.**
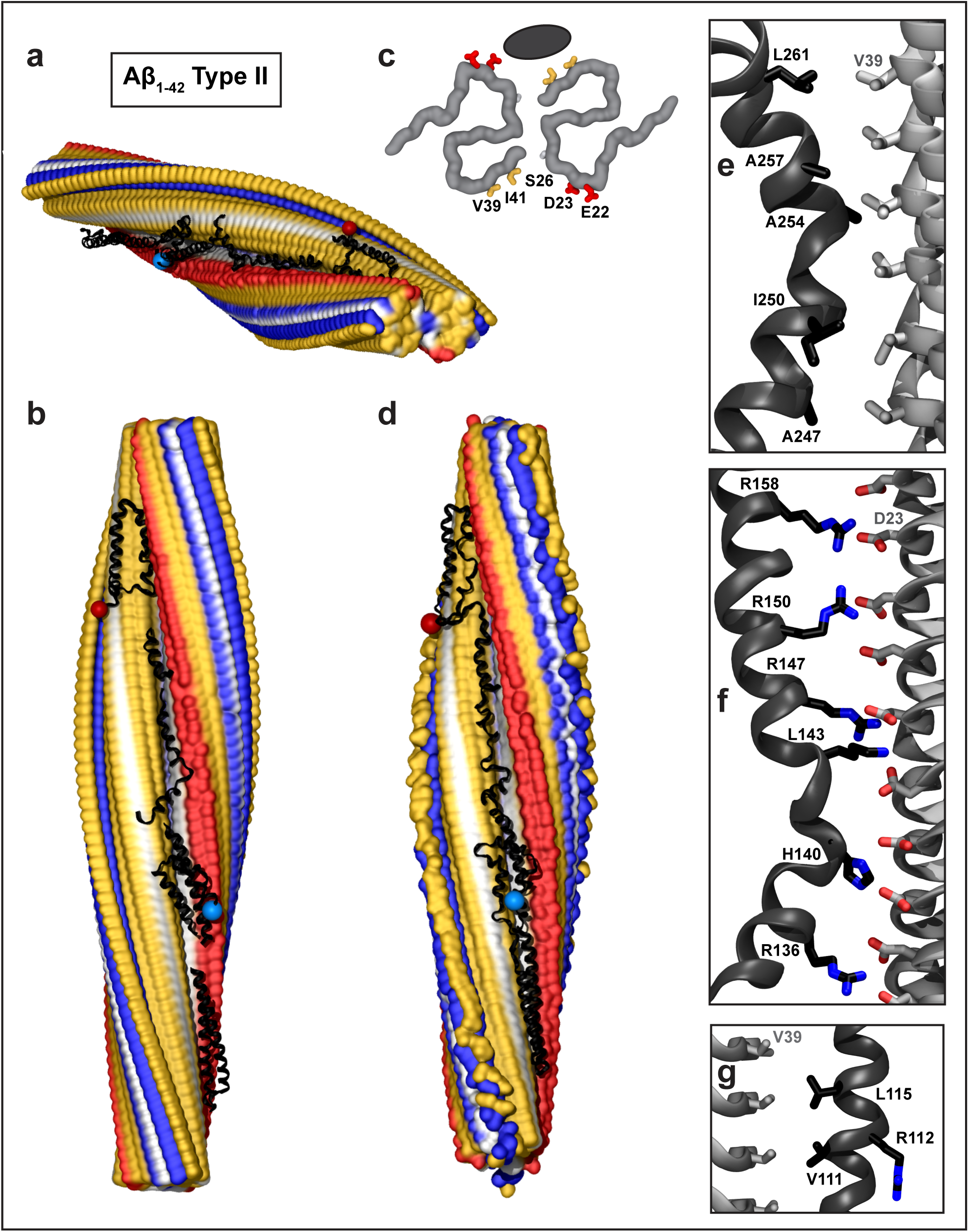
Structural model of full-length apoE in complex with A*β*_1-42_ type II fibril from AD vasculature. (**a, b**) Top and side views of the docking model, which was obtained by using apoE segments shown in supplemental Fig. 2d-h and the fibril structure (PDB ID: 7Q4M). Segment positions are compatible with full-length apoE. (**c**) Main chains of one fibril rung, top view. A black oval marks the apoE docking site at one of the two predicted symmetry-related sites per paired protofilament. ApoE-coordinating residues V39, I41 from molecule 1 and S26, D23, E22 from molecule 2 of the A*β*_1-42_ protofilament are shown. (**d**) The model of the full-length apoE in complex with the A*β*_1-42_ fibril after MD simulations. Panels e-g show apoE-amyloid contacts in selected regions of this model in a representative frame (apoE – black, Aβ – gray). (**e**) Hydrophobic interactions between CTD residues 247-261 of apoE and V39 ladder of A*β*_1-42_. (**f**) R158 in apoE3/E4 and other basic residues in helix 4 form favorable ionic interactions with the D23 ladder in A*β*_1-42_ amyloid. (**g**) Hydrophobic residues of apoE helix 3 interact with residue ladder V39 in Aβ_1-42_ amyloid. ApoE side chain 112 (R112 in apoE4) points away from amyloid.

To probe the stability of this docking model, MD simulations were performed. The starting model contained 56 or 64 fibril rungs and five non-overlapping apoE fragments selected from the highest-scoring Cluspro docking poses and arranged along the fibril length N to C with appropriate spacing, as shown in Fig. 2a, b. This model was solvated, charge-neutralized, ionized and simulated at 300 K for 100 ns as described in Methods. To check repeatability, MD simulations were performed in triplicate using three different sets of apoE fragments selected from the highest-scoring ClusPro docking poses. Next, the minimized coordinates for two independent fragment sets were stitched together into a full-length apoE molecule docked onto the fibril. These two models were then simulated for 100 ns at 300 K. The details for individual MD runs and the final models are reported in the supplemental Figs. 6, 7 and Table 1. The model stability was indicated by converging root mean square deviations (supplemental Fig. 6) and by substantial buried solvent-accessible surface area (13,000-14,000 Å^2^ for full-length apoE, slightly smaller for unconnected fragments), which persisted during the simulations, indicating substantial contribution of the hydrophobic effect to the complex formation (supplemental Fig. 7). These MD results suggest that the proposed models of apoE-A*β* fibril complexes are stable and hence, are physically plausible.

A representative final model containing full-length apoE and 64 fibril rungs is shown in Fig. 2d; zoomed-in views show selected regions (Fig. 2e-g). The model confirms that the binding involves mainly hydrophobic interactions (Fig. 2e, g). Additionally, the basic residues of apoE can form favorable ionic interactions with the acidic ladder in A*β*_1-42_ amyloid. Fig. 2f depicts possible salt bridges between the basic residues from apoE helix 4 and D23 ladder in amyloid. Like in Fig. 1, apoE residue 158 projects towards amyloid while residue 112 points away from it (Fig. 2f, g). Unlike Fig. 1, which showed unfavorable interactions between R158 of apoE and M35 of A*β*_1-40_, Fig. 2f suggested favorable interactions including a possible salt bridge between R158 of apoE and D23 of A*β*_1-42_; this interaction would have been less favorable in apoE2 that has C158. Comparison of Figs. 1 and 2 suggests that apoE-fibril interactions depend critically on the structure of the amyloid polymorph and are modulated by the apoE isoforms.

#### ApoE docking to type I Aβ_1-42_ fibrils

Type I A*β*_1-42_ filaments have highly hydrophobic top/bottom surfaces and hydrophilic sides, with only narrow hydrophobic strips alongside each protofilament (supplemental Fig. 1c, Fig. 3a, b). Not surprisingly, full-length apoE, its NTD, CTD, paired helical fragments from the NTD, and fully extended helical fragments from other apoE regions strongly preferred docking at the filament ends (supplemental Figs. 4c, 5c). Still, few docking poses alongside the filament were coordinated by A*β* residue ladders E11, H13, H14, and K16, mainly *via* their hydrophobic moieties. Since the area at the fibril ends was insufficient to accommodate more than one or two apoE fragments, the ends were blocked in the next round of docking, resulting in nearly all docking poses for all apoE fragments associated with the narrow hydrophobic groove along the fibril side. To assemble the docked fragments into a full-length apoE, we selected paired helices 1 and 4 docked at the fibril top (Fig. 3a, b, e, g); these fragments showed strong preference for binding at the top surface (22 out of 30 docking poses, supplemental Fig. 5c). Paired helices 2/3 ran along the narrow hydrophobic strip on one protofilament, and extended helices from the lock and CT regions ran along its counterpart from the other protofilament. Fig. 3a, b shows the proposed model, which is consistent with full-length apoE bound to type I fibril. Zoomed-in views illustrate hydrophobic apoE-A*β*_1-42_ interactions at the fibril top and sides (Fig. 3e, f, h) and the orientation of R158 and R112 apoE side chains that do not interact favorably with amyloid in this model (Fig. 3g, h). Favorable ionic interactions at the fibril side are also possible, including potential salt bridges between K16 ladder of A*β*_1-42_ and acidic residues of apoE helix 2 (Fig. 3i).

**Fig. 3.**
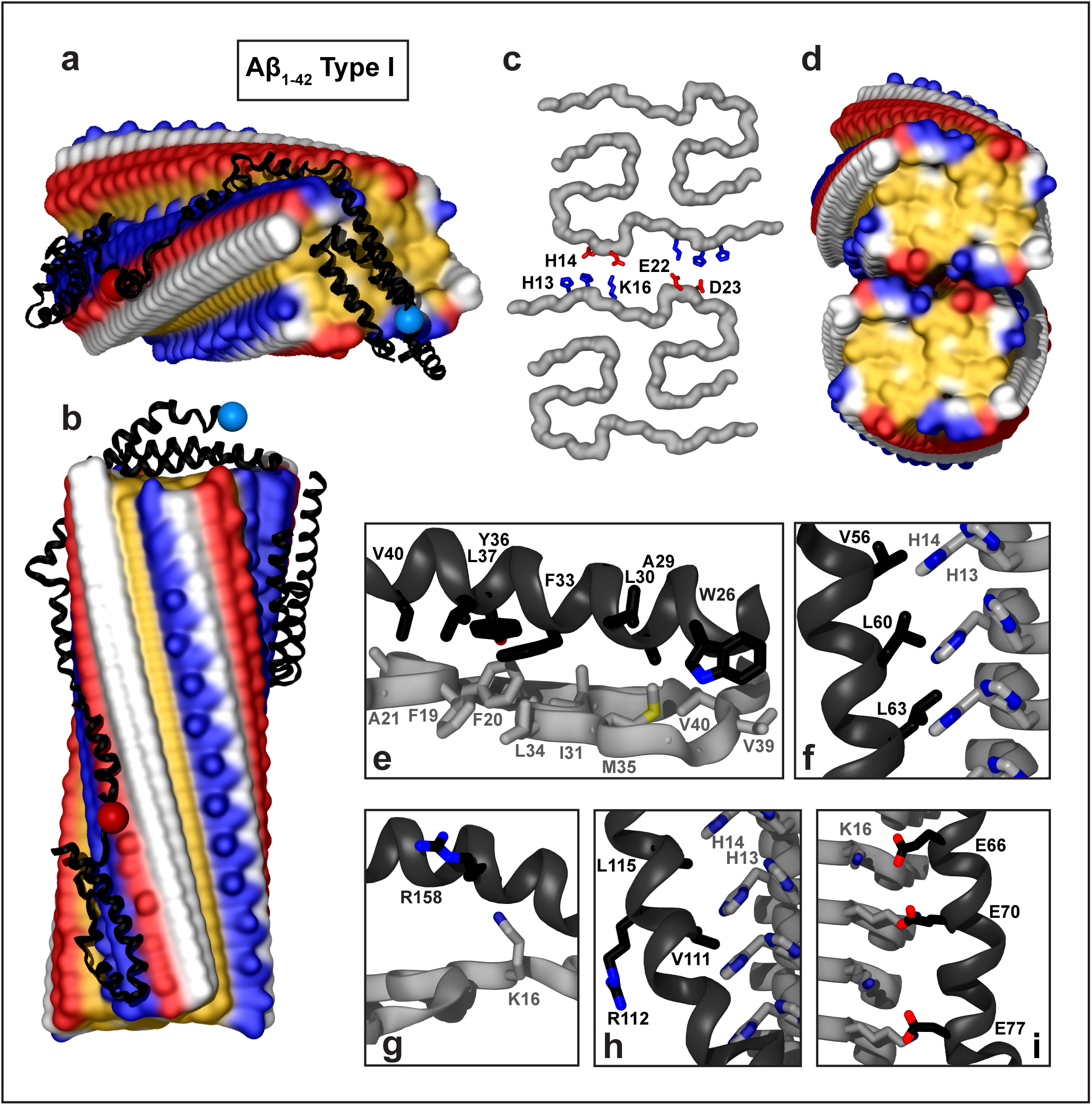
Structural model of apoE docked onto A*β*_1-42_ type I filament from parenchymal deposits. The model was obtained by using apoE segments shown in supplemental Fig. 2d-h and the fibril structure (PDB ID: 7Q4B). Segment positions are compatible with full-length apoE. (**a, b**) Top and side views of the docking model (apoE in black ribbon, amyloid fibril in surface representation). (**c**) One rung of the dimeric type Ib Aβ_1-42_ filament main chain (top view). Residues H13, H14, K16, E22, and D23 forming salt bridges between the two protofilaments [15] are shown. (**d**) Hydrophobic surface representation of type Ib dimeric filament (top view); this hydrophobic fibril end is predicted to form the preferred apolipoprotein binding site. Panels e-i show apoE-amyloid contacts within ∼5Å in selected regions (apoE – black, Aβ – gray). (**e**) Hydrophobic interactions between helix 1 of apoE and the top surface of the A*β*_1-42_ type I filament. (**f**) Hydrophobic residues of apoE helix 2 pack alongside H13, H14 ladder of the A*β*_1-42_ filament. (**g**) R158 in helix 4 of apoE3/E4 does not form favorable interactions at the fibril top. (**h**) Hydrophobic residues of apoE helix 3 interact with hydrophobic residue ladders H13 and H14 in Aβ_1-42_ filament. Side chain 112 (R112 in apoE4) points away from amyloid. (**i**) Acidic residues of apoE helix 2 form electrostatic interactions with the K16 ladder of A*β*_1-42_.

Similar models could be envisioned for the dimeric type Ib filament (Fig. 3c, d). In one model, paired helices 1/4 and 2/3 from NTD of apoE bind at the hydrophobic ends of the two adjacent A*β*_1-42_ protofilaments, while the extended CTD helices bind in the hydrophobic groove along the fibril side. Alternatively, CTD helices, which are highly hydrophobic, may bind across the hydrophobic fibril ends, while the autonomously folded NTD may either remain globular or bind along the fibril side in a partially or fully extended conformation. Such apoE binding is expected to hamper the filament elongation and dissolution and bridge two protofilaments together *via* their ends.

### ApoE docking to the structure of Aβ_1-40_ fibrils grown from the AD tissue-derived seed

The structure of the seeded fibril, which was determined by solid state NMR, is trimeric, with A*β* residues 1-40 forming the amyloid core. A central channel interrupts the hydrophobic end surfaces. The content of this hydrophobic channel is unknown, precluding accurate docking predictions at the fibril ends. The side surfaces in each protofilament show narrow hydrophobic grooves near the A*β* N-terminus, one of which forms the preferred predicted docking site for all apoE fragments (supplemental Fig. 5d). ApoE at this site is coordinated by A*β*_1-40_ residues H6, S8, Y10, V12, H14, with E3 and K16 at the periphery (Fig. 4c). A central ladder comprised of Y10 and V12 flanked by histidines and other A*β*_1-40_ residues forms predominantly van der Waals interactions with the hydrophobic helical faces of apoE (Fig. 4d, g). Additionally, favorable ionic interactions involving apoE helices (e.g. helices 2 and 4, Fig. 4f, e) can contribute to binding. This includes potential salt bridges between an acidic ladder formed by E3 from A*β*_1-40_ and basic residues from helix 4 of apoE (Fig. 4e). Notably, R158 forms such favorable interactions, which are expected to be less favorable for C158 in apoE2. Conversely, residue 112 (R112 in apoE4) projects away from amyloid in this (Fig. 4g) and all other apoE-fibril complexes.

**Fig. 4.**
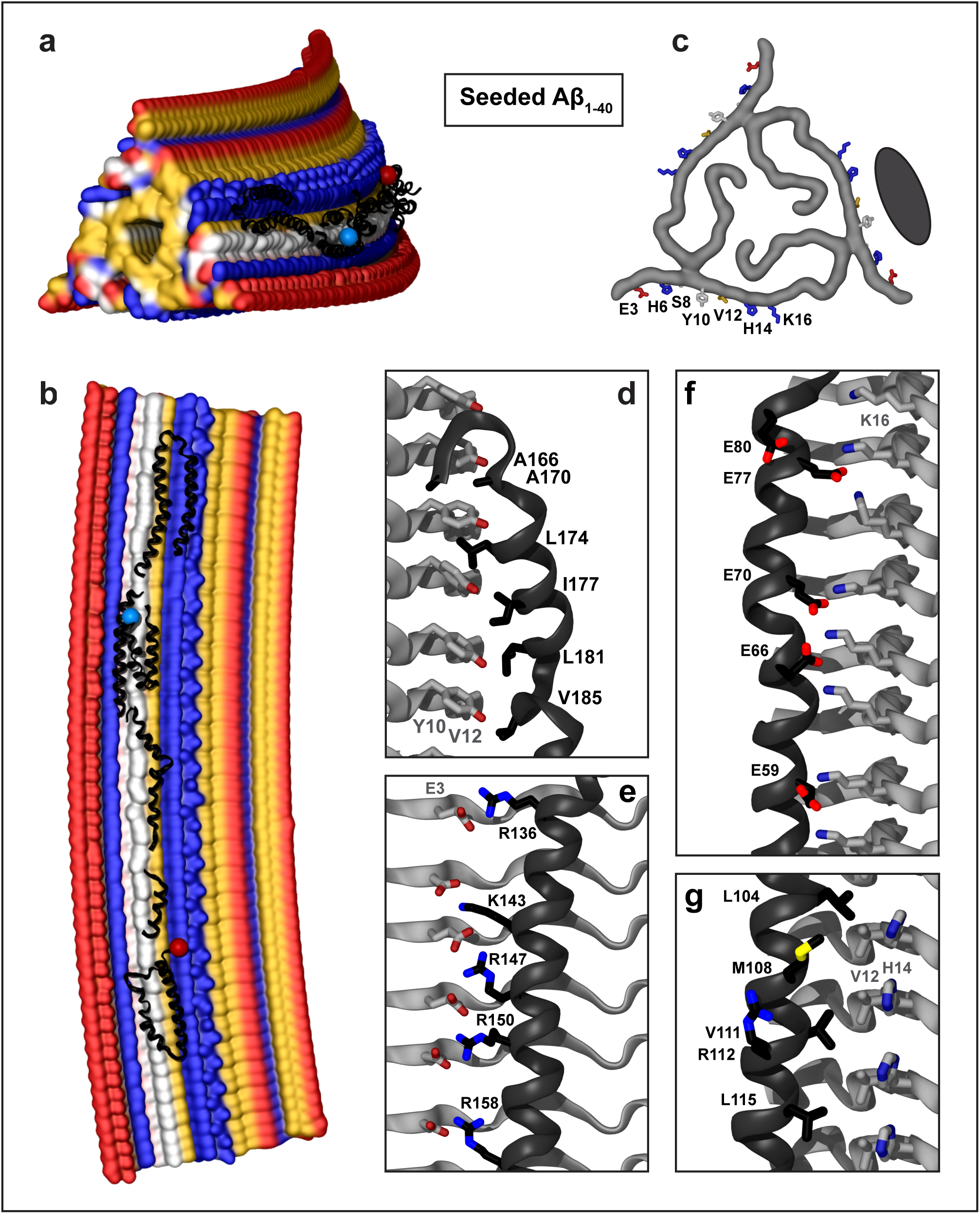
Structural model of apoE docked onto Aβ_1-40_ fibrils obtained by seeding using AD tissue-derived seed. The model was obtained by using apoE segments shown in supplemental Fig. 2d-h and the fibril structure (PDB ID: 2M4J). Segment positions are compatible with full-length apoE. (**a, b**) Top and side views of the docking model with apoE in black ribbon and amyloid fibril in surface representation. (**c**) Top view of one filament layer main chains. ApoE-coordinating residues E3, H6, S8, Y10, V12, H14, and K16 are indicated. Black oval indicates docked apoE. Panels d-f show apoE-amyloid contacts within ∼5Å in selected regions (apoE – black, Aβ – gray). (**d**) Y10, V12 ladder in Aβ_1-40_ interacts with the hydrophobic helical face from the apoE “lock” region. (**e**) E3 ladder in Aβ_1-40_ forms favorable ionic interactions including potential salt bridges with basic residues from apoE helix 4. (**f**) K16 ladder in Aβ_1-40_ forms favorable ionic interactions including potential salt bridges with acidic residues from apoE helix 2. (**g**) Hydrophobic residues of apoE helix 3 interact with residue ladders V12 and H14 in Aβ_1-40_. Side chain of 112 (R112 in apoE4) projects away from amyloid.

**Fig. 5.**
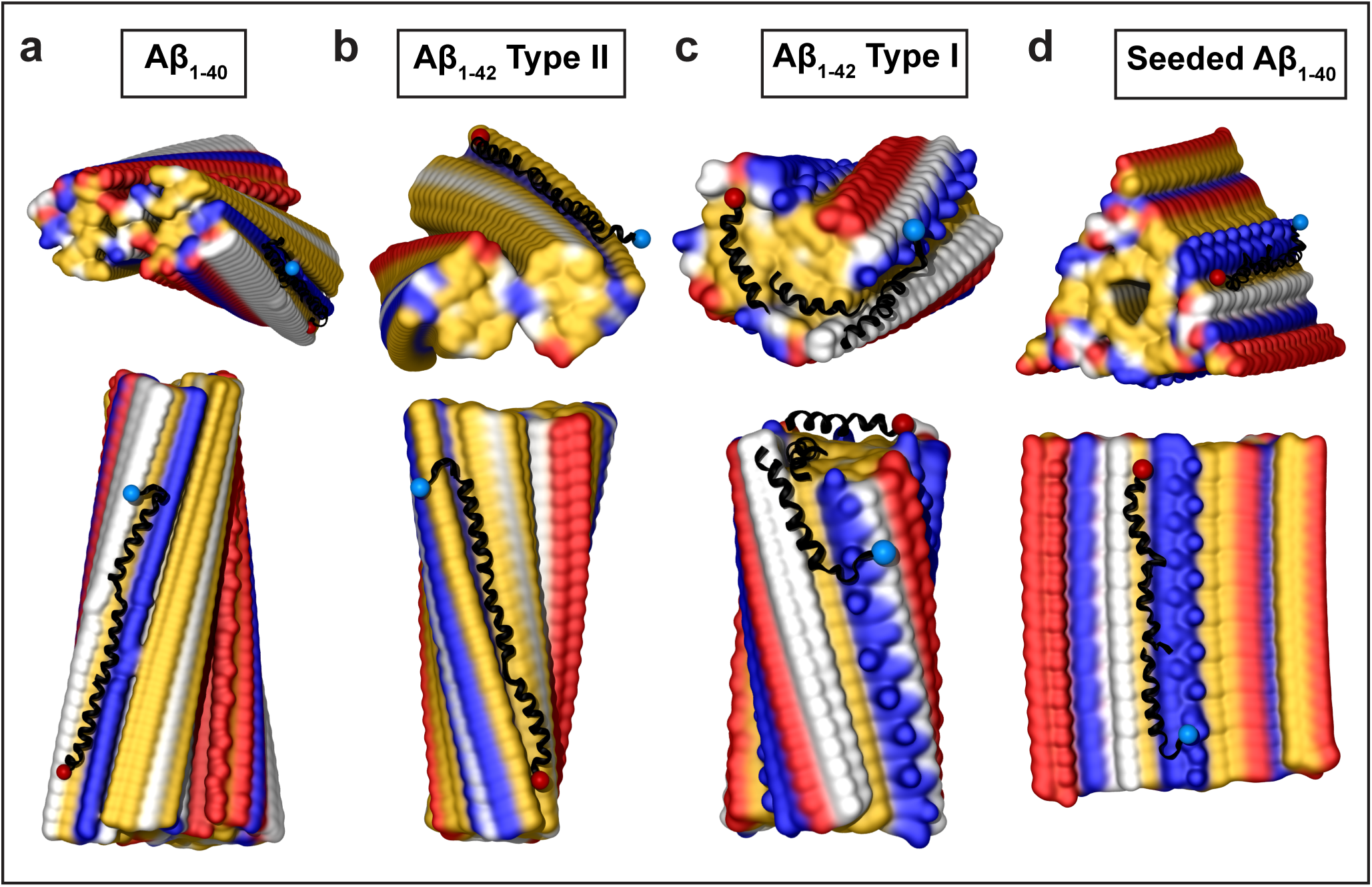
Structural models of apoC-III docked onto A*β* fibrils. The models were obtained by using apoC-III segments shown in supplemental Fig. 3b and the fibril structures of Aβ_1-40_ (a, d) and Aβ_1-42_ (b, c), PDB ID: 6SHS (**a**), 7Q4M (**b**), 7Q4B (**c**), and 2M4J (**d**). Blue and red dots indicate N– and C-termini of docked apoC-III. Segment positions are compatible with full-length apoC-III.

### ApoC-III docking to the structures of Aβ_1-40_ and Aβ_1-42_ fibrils

In addition to lipid-free apoE, the NMR structure of apoC-III was docked to the structures of A*β*_1-40_ and A*β*_1-42_ fibrils. The apoC-III molecule in this structure wraps around the SDS micelle, with the hydrophobic *α*-helical faces projecting inward, mimicking the lipid-bound conformation; relative helical positions are constrained by the micelle size of ∼4.4 nm [65]. Like apoE, full-length apoC-III did not produce any plausible docking poses. Next, apoC-III was split into several highly *α*-helical fragments observed in the NMR structure. Like apoE, all apoC-III fragments docked *via* their hydrophobic helical faces to all amyloids. Consistent docking poses were obtained using three fragments containing residues 1-28, 29-45, and 46-66; partially disordered CT residues 67-79 were excluded. Fig. 5 shows selected docking poses for fragments combined into a model consistent with full-length apoC-III; all docking poses are summarized in supplemental Fig. 8. Like apoE, apoC-III preferentially docked *via* van der Waals interactions to the hydrophobic fibril surfaces; the binding surfaces predicted for apoC-III and apoE substantially overlapped.

In summary, we used the structures of human lipid-free apoE and lipid-bound apoC-III as starting models for docking to four different human A*β* fibrils from patient-based amyloid deposits. The results consistently showed that apolipoprotein-fibril binding is driven by hydrophobic interactions, depends upon the structure of the amyloid fibril polymorph, and involves conformational changes in flexible apolipoprotein molecules. For each amyloid fibril, the best docking poses for the two apolipoproteins were similar, validating our approach and suggesting that other apolipoproteins may interact with fibrils in a similar manner.

### Implications for fibril stability and experimental support for the proposed models

#### Apolipoprotein binding may stabilize amyloid filaments

Figures 1-4 illustrate potential modes of apoE-amyloid fibril interactions. These figures show CTD helices fully extended and NTD helices partially extended. Alternative modes may involve various extents of NTD opening, from fully extended to globular; opposite N-to-C orientation of apolipoprotein molecule on the fibril, etc. Apolipoprotein binding is expected to stabilize the filaments by shielding their exposed hydrophobic surfaces from solvent with amphipathic *α*-helices. Amyloid stabilization may also result from apolipoprotein binding between nearby protofilaments. In fact, the predicted binding sites are compatible with protofilament self-association. One example is A*β*_1-40_ fibril morphologies II and III wherein dimeric or trimeric protofilaments pack side by side *via* ionic interactions, leaving large hydrophobic surface sides available for binding apoE (Fig. 1a). Another example is A*β*_1-42_ fibril type Ib where two filaments pack side by side *via* ionic interactions to form a dimer (Fig. 3c, d). Although such dimerization partially occludes the predicted lateral apoE binding site, it permits binding at the hydrophobic filament ends, which are the preferred apoE docking sites in these fibrils (supplemental Fig. 5c). ApoE binding at the ends of two adjacent filaments will likely bridge them together. A similar mechanism could perhaps explain apoE-mediated bridging of apoC-II amyloid filaments reported in cross-linking studies, which required both apoE domains [36]; this mechanism may extend to other apoE-amyloid interactions.

#### Experimental evidence from prior studies

Amyloid fibril structures often feature large hydrophobic surfaces [11], which are probably sequestered *in situ* by amphipathic molecules such as apolipoproteins and lipids that co-deposit with amyloid. These molecules are stripped by detergent during fibril isolation from tissues and hence, are not observed in the structures of tissue-derived amyloids. However, cryo-EM structures of recombinant α-synuclein fibrils grown in the presence of lipids have shown lipid bicelles bound *via* the largely hydrophobic groves along the fibril side [18]. Similar fibril sites are expected to bind other amphipathic molecules such as apoE.

Our predicted apoE binding sites correspond to faint extra densities seen in cryo-EM maps near H13 and H14 of A*β*_1-42_ type I fibrils and near V39 and I41 in A*β*_1-42_ type II fibrils [15]. Such densities are consistent with apoE binding but do not prove it, necessitating additional experimental verification. Our proposed mechanism involves conformational changes in apoE to expose its hydrophobic helical faces for binding alongside amyloid fibrils or at their ends. This mechanism can be extrapolated to other fibrils that also have exposed hydrophobic surfaces. Therefore, prior experimental studies of apoE-A*β* amyloid interactions can provide evidence for our proposed binding mechanism, even though the amyloid structures in those studies are unknown and can differ from the structures of tissue-derived fibrils used in the current work.

The first supporting evidence comes from early EM studies of negatively stained A*β*_1-28_ and A*β*_1-40_ fibrils grown in the presence of apoE that was visualized using immunogold-labeled antibodies to its NT, CT and central parts [32]. ApoE bound along the whole fibril length with all apolipoprotein parts detected on the fibrils, supporting apoE binding along the fibril side suggested in our models (Figs. 1-4).

Second, binding assays consistently showed that *β*-sheet formation and aggregation of A*β* increase its affinity for apoE [33] and reported that high-avidity binding requires A*β* residues 12-28 [38], which are in the cores of the patient-derived fibrils [15, 63, 64]. Collectively, these findings indicate that A*β* fibril formation facilitates high-affinity binding of apoE, in excellent agreement with our models.

Third, apoE fragments co-deposit with A*β* in human AD brain [40–42], justifying our fragment-based approach. Moreover, *in vitro* studies of apoE fragments reported that all apolipoprotein parts, including NT, hinge, lock, and CT regions, bind A*β* amyloid with generally lower affinity than the full-length apoE [32, 33, 35, 36, 47, 55, 57]. FRET studies of lipid-free apoE indicated that NTD opens upon binding to A*β*, and cross-linking studies reported increased distance between the NT helices h1 and h3 upon binding to A*β* fibrils [35, 36, 47, 55, 57]. Collectively, these findings strongly support domain separation and NTD helix bundle opening in apoE upon binding to A*β* amyloid, which is a key feature of our models.

Fourth, cross-linking and mass spectrometry studies of full-length apoE have mapped the A*β* monomer binding site onto the hydrophobic interface between NTD and CTD [34], while fluorescence labeling studies of CTD reported an overlap between the binding sites for lipids and A*β* [73]. Consequently, hydrophobic helical faces from both NTD and CTD domains of apoE bind A*β*.

Fifth, binding of sub-stoichiometric amounts of apoE (∼1:100 apoE:A*β*) was reported to stabilize A*β* fibrils ([53, 55, 56, 62] and references therein). This ratio is consistent with our lateral binding models where one apoE molecule spans 50-60 fibril layers (∼90 layers in a fully extended conformation) for an approximate binding stoichiometry of 1:90 apoE:A*β*.

Sixth, many kinetic studies reported that at sub-micromolar concentrations and sub-stoichiometric ratios (∼1:100 apoE:A*β)*, apoE delays A*β* fibril formation [35, 52, 53, 56, 59–62]. In these studies, apoE extended the lag phase in the unseeded fibrillation, suggesting that apoE binds and stabilizes A*β* multimers and short fibrils formed during the lag phase [52, 53, 60, 61]. ApoE was also reported to decelerate fibril elongation and maturation in most [52, 53, 56, 62] but not all [61] seeded experiments. Taken together with our models, these results suggest that, depending on the structure of the amyloid polymorph, apoE can bind along its side (to hamper fibril fragmentation and secondary nucleation) and/or at its ends (to block amyloid maturation and elongation). Therefore, our models suggest that the effects of apoE on amyloid growth and proliferation are amyloid polymorph-specific, which helps reconcile controversial findings from prior studies.

Lastly, our proposed mechanism of apoE-amyloid interactions resembles the mechanism of apoE-lipid interactions determined by using structural NMR, cross-linking/mass spectrometry, FRET, EPR and other methods [27, 28, 31, 34, 74]. This similarity suggests a unified molecular mechanism for apoE binding to extended hydrophobic surfaces in amyloids and lipids. This idea is supported by experimental studies reporting that lipids and A*β* amyloid oligomers bind at the same site of apoE CTD [73]. Additionally, surface plasmon resonance and competitive binding studies concluded that the same site in apoE binds to various amyloids, including A*β* [75]. Together, these findings support a unified molecular mechanism of apoE binding to various amyloids. We propose that this mechanism resembles apoE-lipid binding.

## DISCUSSION

This study harnessed cryo-EM structures of tissue-derived and seeded human A*β*_1-40_ and A*β*_1-42_ fibrils [15, 63, 64] to propose molecular models of full-length apoE bound to A*β* amyloids. Key aspects of these models have been supported by extensive experimental evidence from prior studies. The stability of a selected model of apoE-A*β*_1-42_ fibrils was verified in MD simulations (Fig. 2). Additional support comes from apoC-III docking to these fibrils (Fig. 5). As apolipoprotein starting models we used either lipid-free apoE (34 kDa) or lipid-bound apoC-III (9 kDa), yet the predicted amyloid-bound conformations were strikingly similar (Figs. 1-4 vs. 5). Since our approach yields consistent results for two different apolipoproteins in various initial conformations docked to four different fibril polymorphs, it probably extends to other apolipoproteins and other amyloids.

### ApoE binds lipids and amyloids *via* similar molecular mechanisms

The models in Figs. 1–4 exemplify how apoE can interact with various disease-relevant A*β* fibrils. Alternative models may include various degrees of NTD opening, opposite N-to-C orientation of apoE on the fibril, one apoE molecule binding at the ends of two or more adjacent protofilaments, etc. These models exemplify the general mechanism of apolipoprotein-amyloid fibril binding: i) hydrophobic surfaces, which are exposed along the side of amyloid filaments or at their ends, can interact favorably with hydrophobic faces of amphipathic apolipoprotein *α*-helices; ii) apolipoprotein molecules undergo major conformational changes to expose these hydrophobic helical faces for ligand binding. Importantly, this mechanism can be extrapolated to apoC-III (Fig. 5) and other apolipoprotein-amyloid interactions. Moreover, it resembles the binding of exchangeable apolipoproteins to lipids, which is necessary for normal lipoprotein metabolism [28]. Therefore, we propose that both lipid transport and amyloid binding by apoE and other apolipoproteins (apoAs, apoCs) that co-deposit with amyloids involve similar molecular mechanisms. These mechanisms utilize large hydrophobic faces of flexibly linked apolipoprotein *α*-helices for binding to hydrophobic surfaces in lipoproteins and in amyloids.

### ApoE may act as a pro– or anti-amyloid chaperonin with cryptic sites

The proposed mechanism of apoE binding to amyloid can be compared to the action of Brichos domain, an anti-amyloid chaperonin whose hydrophobic amyloid binding site is occluded in the apoprotein by a dynamic helical segment [76]. Brichos binding along the hydrophobic side to A*β*_1-42_ fibrils blocks secondary nucleation, and thereby delays amyloid proliferation [77]. We speculate that lateral binding of apoE at the hydrophobic fibril side has a similar effect. Conversely, capping the fibril ends will likely block their elongation and thereby shift the A*β* aggregation towards secondary nucleation [77], which is proposed to be the major mechanism of A*β* amyloid proliferation [78]. Therefore, we posit that, depending on the structure of the amyloid polymorph and the apoE:A*β* ratio, apoE may act as either an anti– or a pro-amyloid chaperonin: preferential binding at the fibril side will likely hinder amyloid proliferation *via* secondary nucleation, while capping the fibril ends may have an opposite effect. These polymorph-dependent effects help reconcile prior reports on the pro– or anti-amyloid action of apoE [8, 53].

Amyloid binding by apoE and Brichos can be compared with ligand binding to cryptic allosteric sites in globular proteins. Cryptic sites are occluded in ligand-free proteins. These sites tend to have a strong binding hot spot that can be identified computationally; they exhibit high flexibility in the adjacent regions, which facilitates conformational changes necessary for ligand binding; moreover, ligand binding at cryptic sites has elements of induced fit [72]. We suggest that in apolipoproteins, the binding hot spots comprise hydrophobic helical faces, high flexibility is conferred by interhelical linkers, and induced fit enables apolipoproteins to bind to diverse amyloids as well as lipids. Apolipoprotein properties that facilitate such binding are described below.

### Apolipoprotein α-helices form an amyloid and lipid binding motif

Preferential docking of different fragments of apoE and apoC-III to the same fibril sites suggests that the binding is driven by common properties of apolipoprotein *α*-helices. Amino acid sequences of exchangeable apolipoproteins are enriched in hydrophobic residues and contain 11/22-mer tandem sequence repeats with high propensity to form distinct amphipathic *α*-helices [24]. Hydrophobic faces in apolipoprotein *α*-helices span 30-50% of helical circumference and are nearly straight, with only a small right-handed twist, as compared with a much narrower hydrophobic face and a much larger left-handed twist typical of 7-mer helical repeats in globular proteins [24, 79]. Moreover, NMR studies of lipoproteins reported that apolipoprotein helices can slightly unwind on the lipid, further straightening their hydrophobic faces [80]. Apolipoprotein *α*-helices can span several 11-mer repeats to form extended hydrophobic faces (∼1 nm wide and tens of nm long), which can embed into the lipid monolayer [24]. We propose that such wide, long, non-twisting hydrophobic helical faces facilitate apolipoprotein binding to amyloid. Our models suggest that apolipoprotein binding is compatible with either right– or left-handed twist seen in cryo-EM structures of A*β*_1-40_ or A*β*_1-42_ fibrils (Figs. 1-3, 5).

Additionally, our models suggest that ionic interactions can contribute to apolipoprotein binding in some but not all amyloids. In apolipoprotein *α*-helices, hydrophobic faces are flanked by basic residues; in the lipid-bound state, hydrophobic faces insert into the lipid monolayer to bind acyl chains, while the flanking basic residues can interact with phospholipid head groups [24]. Our models suggest that basic residue arrays in apolipoprotein *α*-helices can interact favorably with acidic residues in some amyloids to form salt bridges (Figs. 2f, 4e). These interactions differ for different amyloid polymorphs. Furthermore, apoE is an apolipoprotein with the highest affinity for negatively charged surfaces [65]; such surfaces are formed by exposed ladders of E22, D23 in A*β*_1-42_ but not in A*β*_1-40_ fibrils explored in the current study. Such structural and physicochemical differences among different amyloid polymorphs perhaps contribute to the discrepant experimental data on apoE-amyloid interactions.

In summary, tandem 11-mer sequence repeats in apolipoproteins encode for amphipathic *α*-helices whose wide long non-twisting hydrophobic faces flanked by basic residues can bind either lipid surfaces or amyloids. Flexible linkers between these helices help expose their hydrophobic faces for binding to various amyloid structures. These properties of apolipoproteins explain why they are found in all amyloid deposits *in vivo* and provide clinical markers of amyloid [1].

### Isoform-specific effects in apoE-amyloid interactions

Our models suggest how apoE can interact with amyloid in an isoform-specific manner. First, apoE domain separation and helix bundle opening in NTD, which we posit is critical to amyloid binding, depends on the apolipoprotein structural stability that is isoform-specific and is lowest for apoE4 [25, 39].

Second, Arg/Cys158, which flanks the hydrophobic helical face in apoE, can interact directly with the A*β* fibrils in a manner that depends upon the apoE isoform and the amyloid polymorph. In apoE3 and apoE4, R158 can form favorable ionic interactions with acidic residues in some fibrils, as suggested by our models of type II A*β*_1-42_ or seeded A*β*_1-40_ fibrils (Fig. 2f, 4e), but for C158 in apoE2 such local interactions would not be as favorable. Conversely, in A*β*_1-40_ morphology I fibrils, R158 in apoE3 and apoE4 forms an unfavorable interaction with M35 (Fig. 1f), which likely becomes more favorable upon Met oxidation in A*β*, which increases with aging, or for C158 in apoE2. These direct interactions are amyloid polymorph-specific.

Other factors influencing isoform-specific apoE-amyloid interactions probably include the levels of apoE (which are lowest for apoE4) and post-translational modifications (which may interfere with conformational opening and ligand binding). Additional effects may stem from lipids and HS, as these ubiquitous amyloid constituents bind apoE in an isoform-specific manner [19].

### Apolipoprotein binding can modulate biological properties of amyloids

We posit that apoE binding to mature amyloid fibrils may have either anti– or pro-amyloid effects depending on the binding mode (lateral or end-capping) and the apoE:A*β* ratio. Additionally, apoE-A*β* interactions during amyloid formation may alter the fibril morphology. Moreover, apolipoprotein binding to amyloid oligomers is expected to stabilize their hydrophobic surfaces and hinder their maturation into fibrils, consistent with reports of apoE delaying A*β* amyloid nucleation and maturation [52, 53, 55, 56, 62]. Apolipoproteins may also compete for binding to amyloid with anti-amyloid chaperones such as clusterin and Brichos [53, 76], as well as other chaperones that bind their substrates *via* multiple hydrophobic contacts [81]. Whether the net effect is beneficial or pathologic probably depends on several factors, such as the location and affinity of the apoE binding sites; the stage of amyloid growth; amyloid toxicity; the mechanism of amyloid proliferation (primary or secondary nucleation or fragmentation); local apoE concentration, post-translational modifications, etc. Additional amyloid modulators may involve lipids and HS [82], which bind both apoE and amyloids and influence their interactions in ways that remain to be elucidated.

Notably, unprotected hydrophobic surfaces have been proposed to act as a “danger signal” activating innate immunity [17]. If so, exposed hydrophobic surfaces in amyloids may contribute to amyloid-associated inflammation, including neuroinflammation, the major contributor to the AD pathology that is influenced by apoE [83]. We speculate that apoE binding to exposed hydrophobic surfaces in amyloid can modulate this danger signal.

In summary, apolipoproteins are expected to bind at exposed hydrophobic surfaces in amyloids and modulate amyloid nucleation, elongation, maturation, fragmentation, proliferation, and downstream effects such as inflammation. Whether the effect of binding is anti– or pro-amyloidogenic depends on several factors including the atomic structures of amyloid polymorphs, which can be disease– and organism-specific, e.g. we predict apoE to preferentially cap the ends of type I A*β*_1-42_ fibrils found in human AD (Fig. 3) but bind laterally to type II A*β*_1-42_ fibrils found in other neurodegenerative human diseases and in a mouse model of AD [15] (Fig. 2). We speculate that the downstream biological effects of binding at fibril ends vs. sides are probably very different.

This helps reconcile confounding data from prior studies reporting that apoE is either a pro– or an anti-amyloid chaperonin [3, 8, 22, 45, 53, 84]. The current study proposes a general structural basis for understanding and, ultimately, harnessing these complex effects.

## Supplementary Information

The online version contains supplementary material, including supplemental Figs. 1-8 and Table 1.

## Abbreviations

A*β*: amyloid-*β*
AD: Alzheimer’s disease
apo: apolipoprotein
CT: C-terminal
CTD: C-terminal domain
EM: electron microscopy
FRET: Förster resonance energy transfer
HS: heparan sulfate
MD: molecular dynamics
NMR: nuclear magnetic resonance
NT: N-terminal
NTD: N-terminal domain
PRIBS: parallel in-register intermolecular *β*-sheets

## Acknowledgements

We thank Drs. John E. Straub and Shobini Jayaraman for extremely helpful discussions. NAMD was developed by the Theoretical and Computational Biophysics Group in the Beckman Institute for Advanced Science and Technology at the University of Illinois at Urbana-Champaign.

## Author contribution

EL performed the research and created the graphics. MNN contributed to research. EL, MJR and OG designed the approach and analyzed the data. OG conceptualized the study and wrote the first draft. EL and MJR contributed to manuscript writing and editing. All authors discussed the results and read and approved the manuscript.

## Data availability statement

All models and protocols are included in the main manuscript and the supplement. The PDB files for the final models of apolipoproteins in complex with fibrils are available from the corresponding author upon reasonable request.

## Disclosure statement

All authors have no financial or any other conflicts of interests to disclose.

## Ethics approval and consent to participate

Not applicable (there are no animals and no human subjects in this study).

## Consent for publication

All authors give their consent to publish this non-human subject study.

## Funding

This work was supported by the National Institutes of Health grants RO1 GM067260 (OG), GM135158 (OG and MNN), and T32 DK007201 (EL); MJR was supported by R01 HL036153.

## Notes

### Competing Interest Statement

The authors have declared no competing interest.

