## supplemental figures and table for "Molecular modeling of apoE in complexes with Alzheimer’s amyloid-β fibrils from human brain suggests a structural basis for apolipoprotein co-deposition with amyloids"

**CONTENT:**

**Supplemental Figures 1-8, Supplemental Table 1**

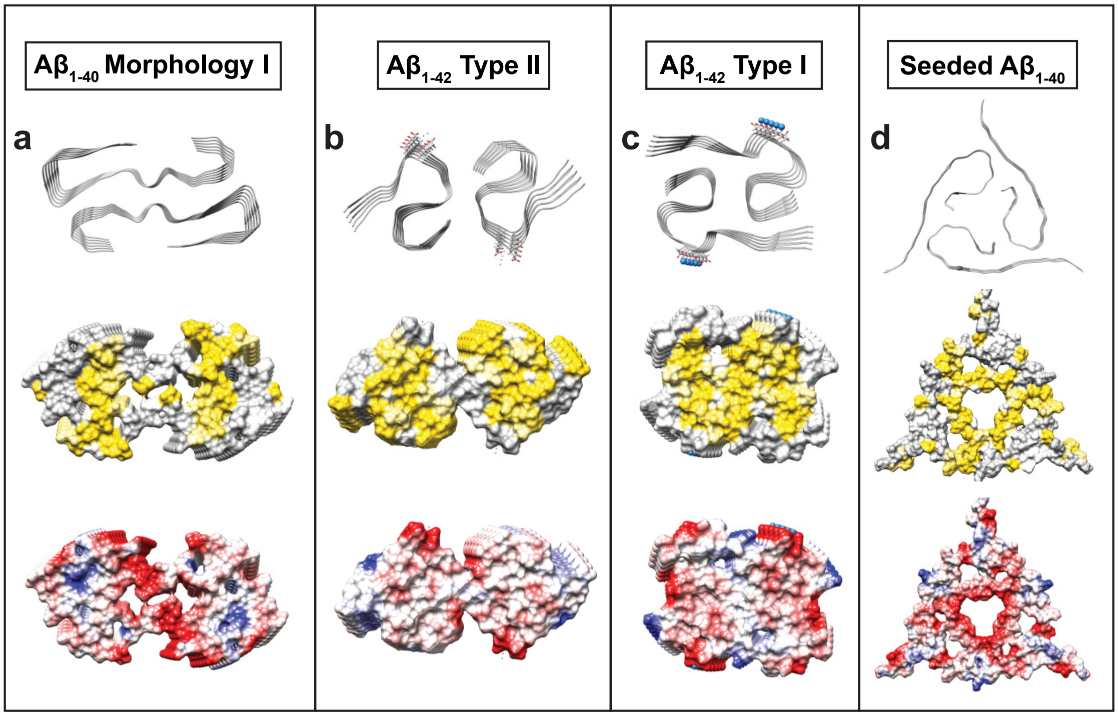


**Supplemental Fig. 1** Structural polymorphs of patient-based $A\beta_{1-40}$ and $A\beta_{1-42}$ fibrils used in the current study. Views down the fibril axis show ribbon diagrams (top) and space-filling models depicting hydrophobic (middle) and Coulombic surfaces (bottom). Color coding in these and in other space-filling models: hydrophobic – yellow, acidic – red, basic – blue.

(**a**) $A\beta_{1-40}$ fibril morphology I from AD vasculature, cryo-EM structure (PDB ID: 6SHS);
(**b**) $A\beta_{1-42}$ type II from parenchymal deposits, cryo-EM structure (PDB ID: 7Q4M);
(**c**) $A\beta_{1-42}$ type I from parenchymal deposits, cryo-EM structure (PDB ID: 7Q4B);
(**d**) $A\beta_{1-40}$ seeded fibril with seed isolated from AD brain tissue, NMR structure (PDB ID: 2M4J).

In panels b and c, small blue spheres indicate putative metal ions bound *via* acidic ladders of E22, D23.

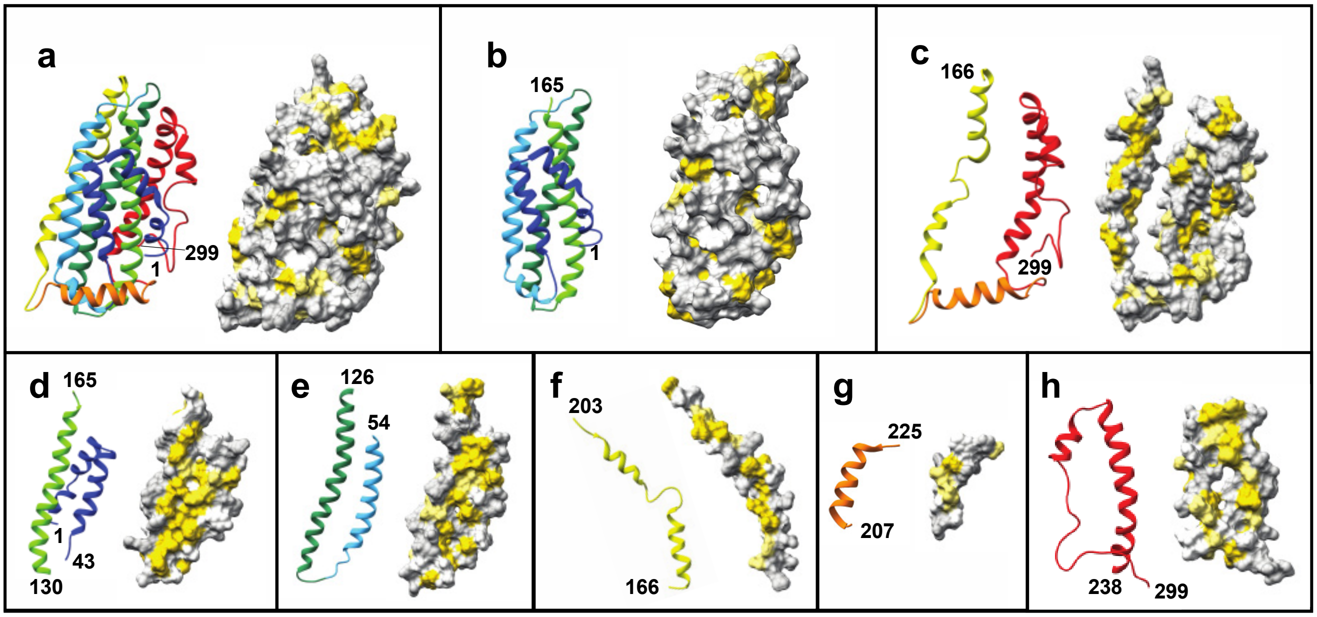


**Supplemental Fig. 2** Solution NMR structure of full-length lipid-free modified human apoE3 (PDB ID: 2L7B) and its fragments used in the current study. Helices in the ribbon diagrams are rainbow-colored N to C (blue to red). Space-filling models are oriented to show hydrophobic surfaces (in yellow). First and last residues in each fragment are shown by numbers.
(**a**) Full-length apoE; (**b**) NTD; (**c**) CTD. (**d-h**) Segments that yielded consistent results in ClusPro docking to amyloid fibrils: (**d**) helices 1 and 4, (**e**) helices 2 and 3, (**f**) hinge, (**g**) lock, and (**h**) CT region.

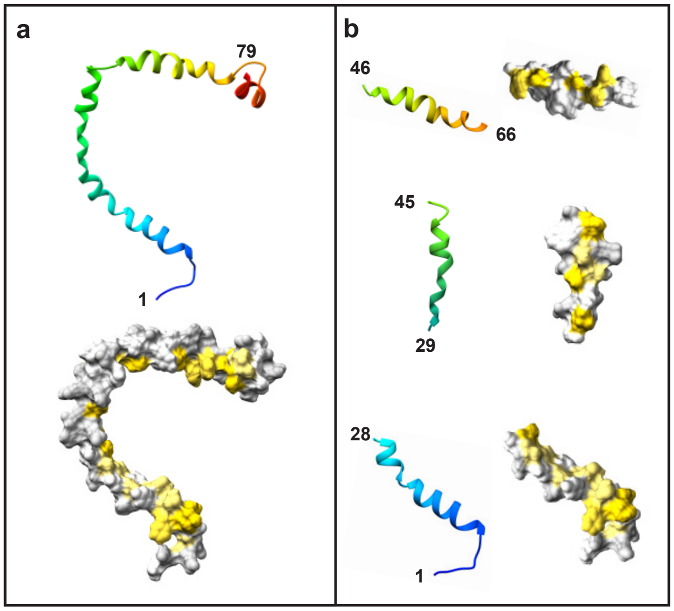


**Supplemental Fig. 3** Solution NMR structure of full-length human apoC-III (PDB ID: 2JQ3) and its fragments used in the current study. The structure, which was determined by using the apolipoprotein bound to SDS micelles (diameter ~4.4 nm), represents the lipid-bound conformation. Ribbon diagrams are rainbow colored N to C (blue to red); space-filling models are oriented to show hydrophobic surfaces (in yellow). First and last residue numbers are indicated. (**a**) Full-length apoC-III. (**b**) NT, middle, and truncated CT fragments, which yielded consistent results in ClusPro docking to amyloid fibrils.

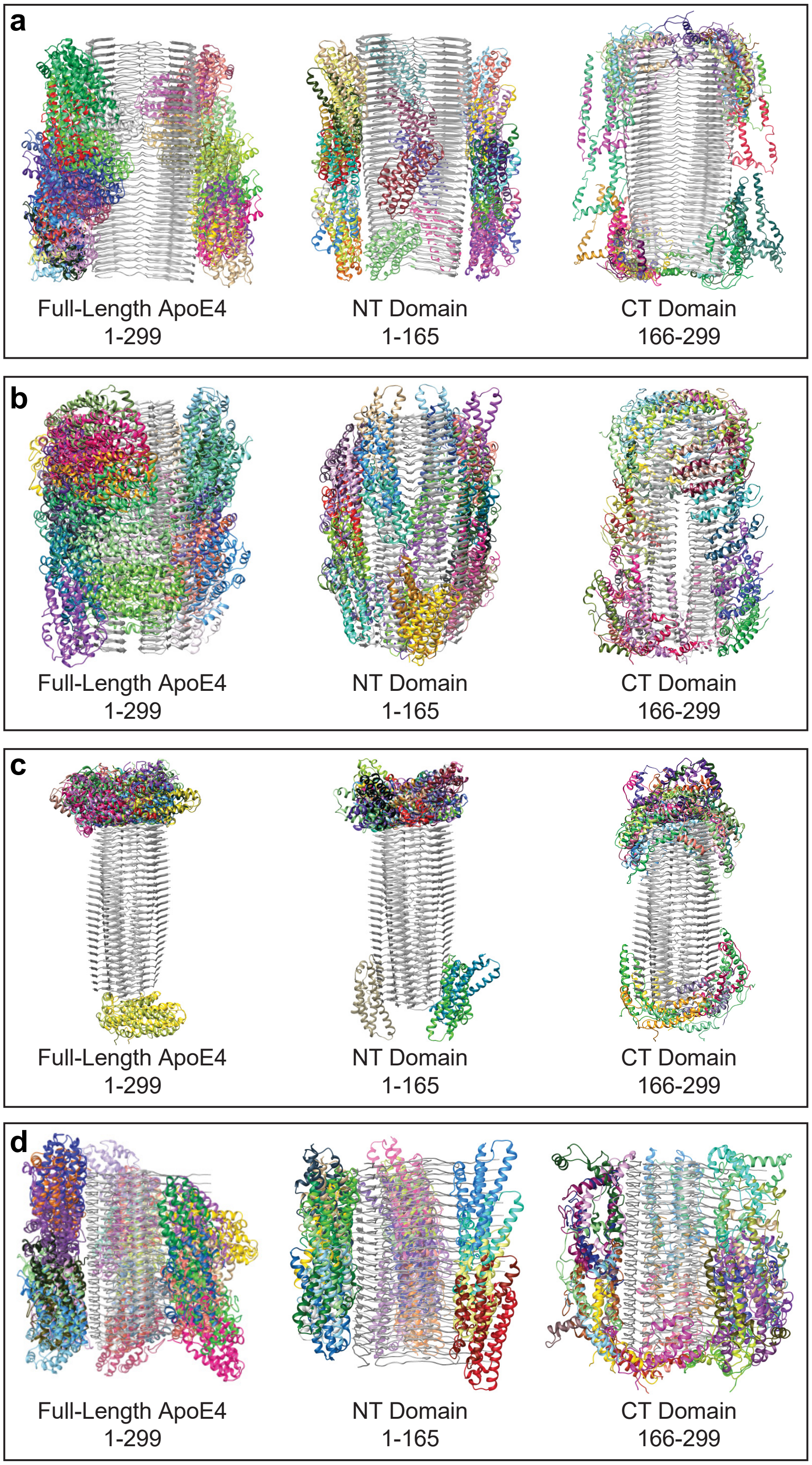


**Supplemental Fig. 4** Summary of docking poses for full-length apoE and its NTD and CTD fragments docked using Cluspro onto four Aβ fibrils. Residues in each fragment are shown by numbers. The top-scoring docking poses for each fragment are shown; each pose is colored differently. The fibrils are:
(**a**) $A\beta_{1\text{-}40}$ fibril morphology I from AD vasculature, cryo-EM structure (PDB ID: 6SHS);
(**b**) $A\beta_{1\text{-}42}$ type II from parenchymal deposits, cryo-EM structure (PDB ID: 7Q4M);
(**c**) $A\beta_{1\text{-}42}$ type I from parenchymal deposits, cryo-EM structure (PDB ID: 7Q4B);
(**d**) $A\beta_{1\text{-}40}$ seeded fibril with the seed isolated from AD brain tissue, NMR structure (PDB ID: 2M4J).

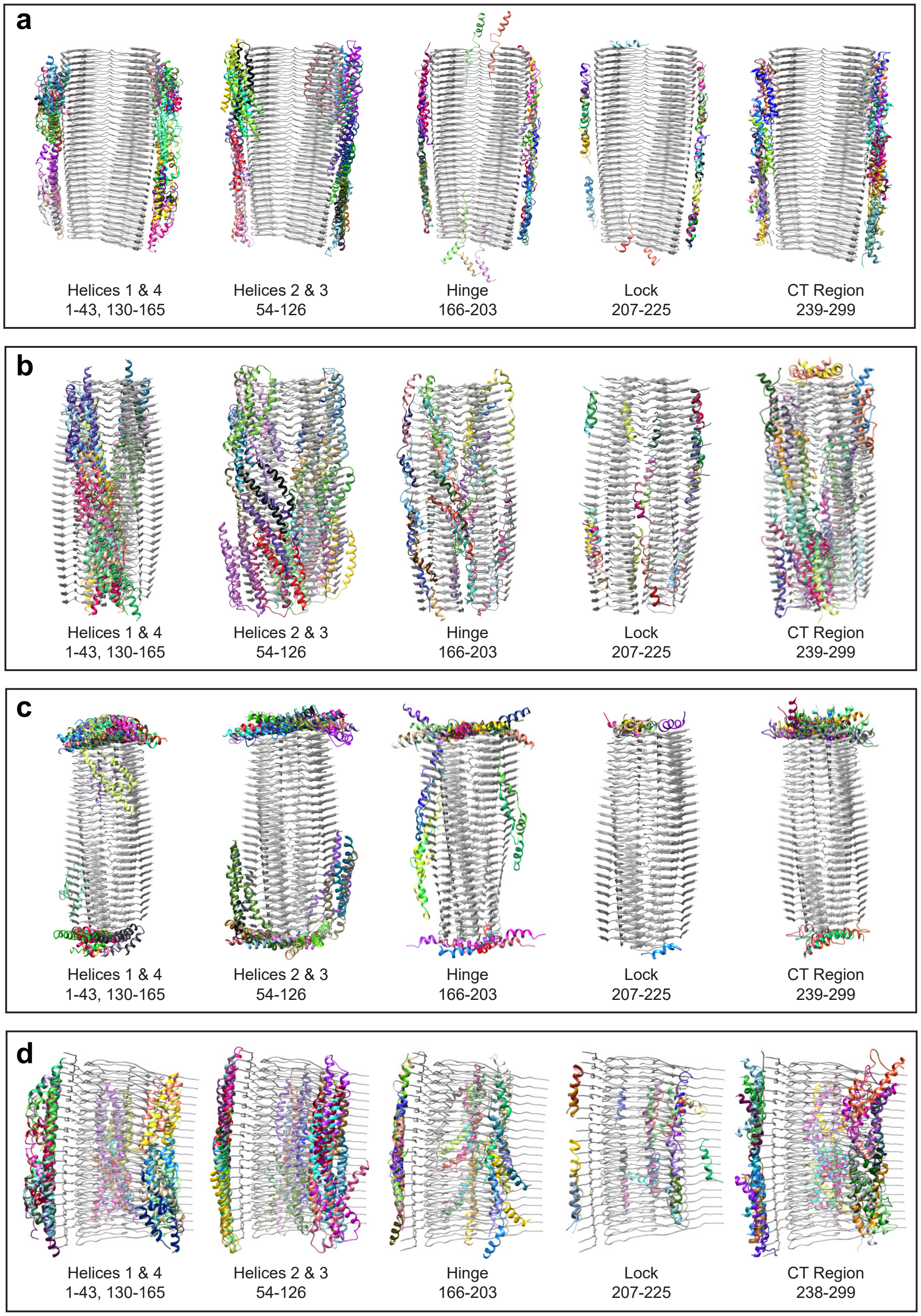


**Supplemental Fig. 5** Summary of docking poses for apoE fragments docked onto four different Aβ fibrils. ApoE was split into fragments shown in supplemental Fig. 2d-h; the residues in each fragment are shown by numbers. The top-scoring docking poses for each fragment are shown; each pose is colored differently. The fibril structure is in gray. The fibrils are:

(**a**) $A\beta_{1\text{-}40}$ fibril morphology I from AD vasculature, cryo-EM structure (PDB ID: 6SHS);

(**b**) $A\beta_{1\text{-}42}$ type II from parenchymal deposits, cryo-EM structure (PDB ID: 7Q4M);

(**c**) $A\beta_{1\text{-}42}$ type I from parenchymal deposits, cryo-EM structure (PDB ID: 7Q4B);

(**d**) $A\beta_{1\text{-}40}$ seeded fibril with seed isolated from AD brain tissue, NMR structure (PDB ID: 2M4J)

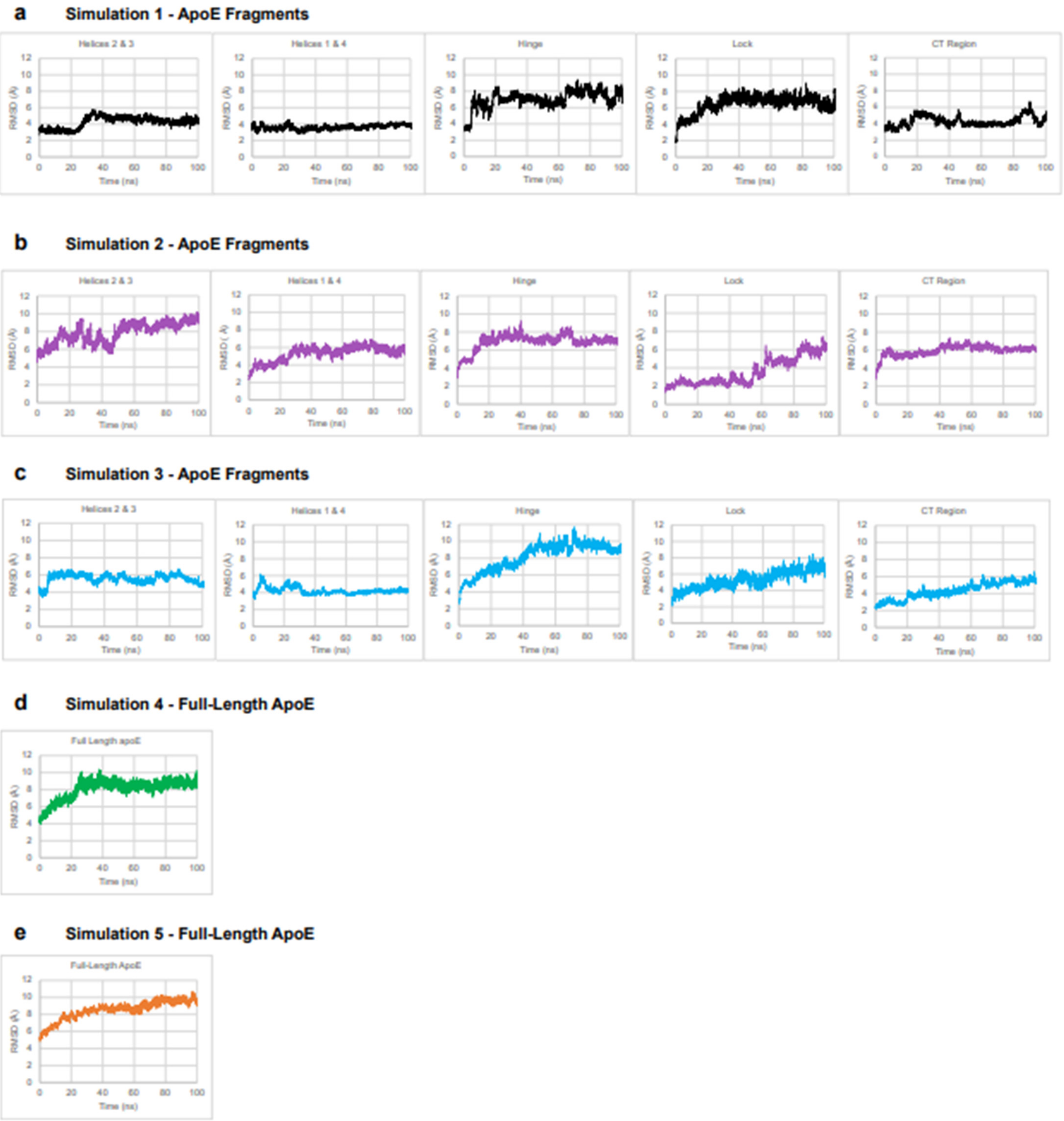


**Supplemental Fig. 6** Root mean square deviations (RMSD) for the backbone atoms in MD simulations of apoE in complex with $A\beta_{1\text{-}42}$ type II fibril during production run time. Each model contained either 56 (a, b, d) or 64 fibril rungs (c, e). RMSD are shown for (**a-c**) triplicate simulations of each apoE fragment, and for (**d, e**) duplicate simulations of the full-length apoE model generated from these fragments as described in the text. For RMSD measurements, each frame was aligned by $C_\alpha$ atoms either by fibril segments adjacent to the fragment of interest (in **a-c**) or the entire fibril (in **d-e**).

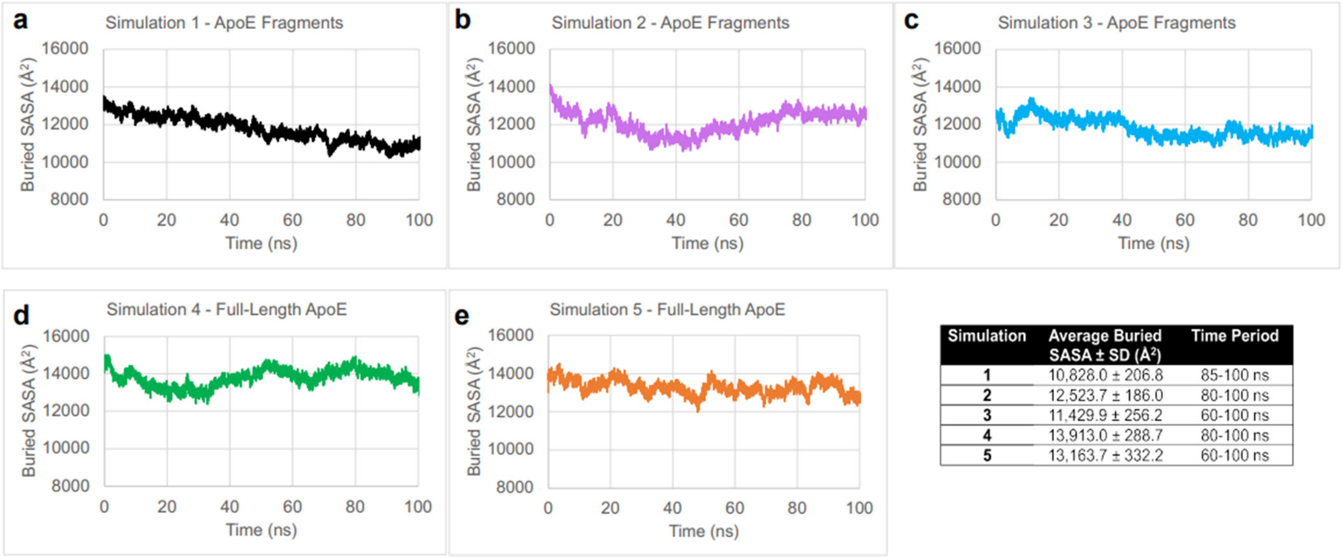


| Simulation | Average Buried SASA ± SD (Å²) | Time Period |
|---|---|---|
| 1 | 10,828.0 ± 206.8 | 85-100 ns |
| 2 | 12,523.7 ± 186.0 | 80-100 ns |
| 3 | 11,429.9 ± 256.2 | 60-100 ns |
| 4 | 13,913.0 ± 288.7 | 80-100 ns |
| 5 | 13,163.7 ± 332.2 | 60-100 ns |

**Supplemental Fig. 7** Buried solvent-accessible surface area (SASA) between apoE and A$\beta_{1\text{-}42}$ fibril during MD simulations. The values are shown for (**a-c**) triplicate simulations of apoE fragments in complex with A$\beta_{1\text{-}42}$ type II fibril, and for (**d, e**) duplicate simulations of the full-length apoE – fibril model generated from these fragments. Respective models are shown in Fig 2b, d of the main manuscript. Average buried SASA for the last 20-40 ns of the production run is tabulated for each simulation.

| *Aβ* | *ApoE* |
|---|---|
| *Glu22* | Lys143 |
| *Asp23* | Lys95, Gln98, Arg136, Ser139, Arg142, Lys143, Lys146, Arg147, Arg150, Arg158, Arg217, Met221, Arg224, Arg274, Gln279, Gln284, Thr289, Ser290 |
| *Val24* | Arg147, Met221, Arg224, Gln284, Thr289, Ser290 |
| *Gly25* | His140, Arg147, Met218, Met221, Arg224, Gln284, Ala285, Thr289, Ser290 |
| *Ser26* | His140, Met218 |
| *Val36* | Arg25, Leu28, Tyr36, Trp39, Gln81, Arg167 |
| *Gly37* | Arg25, Leu28, Ala29, Tyr36, Trp39, Leu63, Gln81, Arg167, Glu168 |
| *Gly38* | Arg25, Trp26, Leu28, Ala29, Tyr36, Trp39, Val56, Leu60, Leu63, Tyr74, Gln81, Glu168, Gly169 |
| *Val39* | Trp26, Ala29, Leu30, Phe33, Tyr36, Leu27, Trp39, Val40, Leu60, Leu63, Met64, Thr67, Leu71, Tyr74, Leu78, Thr83, Glu87, Glu88, Leu104, Met108, Val111, Tyr118, Met125, Leu133, Leu137, Leu159, Tyr162, Gln163, Gly169, Arg172, Gly173, Ala176, Ile177, Leu198, Ala199, Arg215, Ile250, Arg251, Gln253, Ala254, Ala257, Phe265, Val294, Pro295 |
| *Ile41* | Ala91, Arg92, Arg103, Val111, Arg114, Met125, Leu133, Leu137, His140, Leu141, Asp151, Leu155, Leu159, Tyr162, Arg172, Ala176, Arg180, Leu181, Gln187, Arg189, Val190, Ala193, Leu198, Ala199, Trp210, Gly211, Leu214, Arg215, Met218, Glu219, Gly222, Ser223, Ile250, Phe265, Ala269, Gln273, Trp276, Ala292 |

**Supplemental Table 1.** Contact table listing close contacts between the backbone atoms of amino acids from apoE and A$\beta_{1\text{-}42}$ fibrils from parenchymal deposits of AD patients. Contacts within 5Å which are seen in ≥75% of frames are listed for simulation 3 of the model containing apoE fragments and A$\beta_{1\text{-}42}$ type II fibril. The model, which is shown in Fig. 2b of the main manuscript, was subjected to MD simulations as described in the text. Representative simulation 3 was used to determine the contacts; RMSD and buried SASA for simulation 3 are shown in supplemental Figs. 6c and 7c.

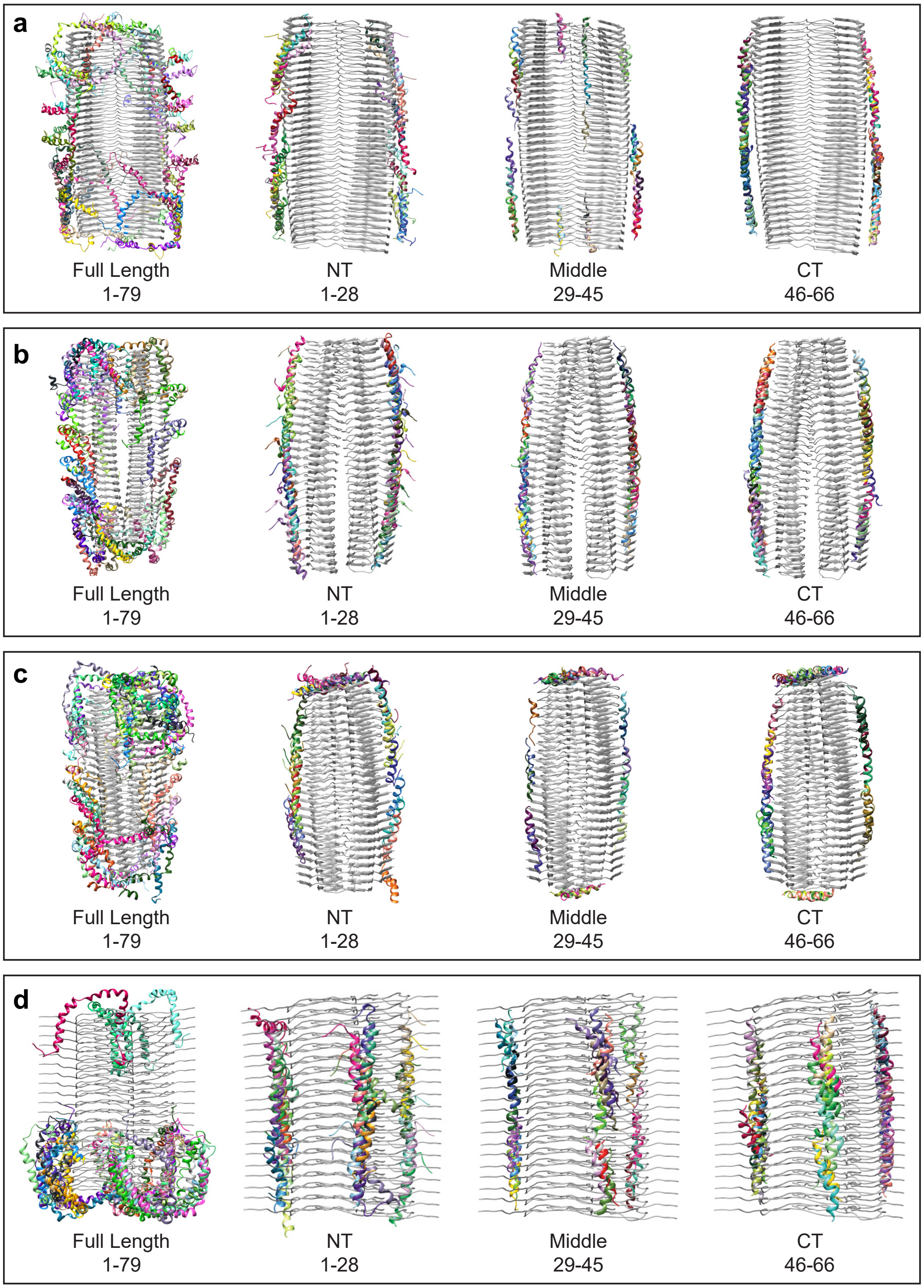


**Supplemental Fig. 8** Summary of docking poses for full-length apoC-III and its NT, middle, and truncated CT fragments docked onto four different Aβ fibrils. Residues numbers in each protein fragment are shown. Top-scoring docking poses are shown, each is colored differently. The fibrils are:
(**a**) $A\beta_{1\text{-}40}$ fibril morphology I from AD vasculature, cryo-EM structure (PDB ID: 6SHS);
(**b**) $A\beta_{1\text{-}42}$ type II from parenchymal deposits, cryo-EM structure (PDB ID: 7Q4M);
(**c**) $A\beta_{1\text{-}42}$ type I from parenchymal deposits, cryo-EM structure (PDB ID: 7Q4B);
(**b**) $A\beta_{1\text{-}40}$ seeded fibril with seed isolated from AD brain tissue, NMR structure (PDB ID: 2M4J).
